# AAV Ablates Neurogenesis in the Adult Murine Hippocampus

**DOI:** 10.1101/2020.01.18.911362

**Authors:** ST Johnston, SL Parylak, S Kim, N Mac, CK Lim, IS Gallina, CW Bloyd, A Newberry, CD Saavedra, O Novák, JT Gonçalves, FH Gage, M Shtrahman

**Author notes:** These authors contributed equally to this work (ST Johnston, SL Parylak). Correspondence to: Fred H. Gage,; Matthew Shtrahman.

## Abstract

Recombinant adeno-associated virus (rAAV) has been widely used as a viral vector across mammalian biology and has been shown to be safe and effective in human gene therapy. We demonstrate that neural progenitor cells (NPCs) and immature dentate granule cells (DGCs) within the adult murine hippocampus are particularly sensitive to rAAV-induced cell death. Cell loss is dose dependent and nearly complete at experimentally relevant viral titers. rAAV-induced cell death is rapid and persistent, with loss of BrdU-labeled cells within 18 hours post-injection and no evidence of recovery of adult neurogenesis at 3 months post-injection. The remaining mature DGCs appear hyperactive 4 weeks post-injection based on immediate early gene expression, consistent with previous studies investigating the effects of attenuating adult neurogenesis. *In vitro* application of AAV or electroporation of AAV2 inverted terminal repeats (ITRs) is sufficient to induce cell death. Efficient transduction of the dentate gyrus (DG)—without ablating adult neurogenesis—can be achieved by injection of rAAV2-retro serotyped virus into CA3. rAAV2-retro results in efficient retrograde labeling of mature DGCs and permits *in vivo* 2-photon calcium imaging of dentate activity while leaving adult neurogenesis intact. These findings expand on recent reports implicating rAAV-linked toxicity in stem cells and other cell types and suggest that future work using rAAV as an experimental tool in the DG and as a gene therapy for diseases of the central nervous system (CNS) should be carefully evaluated.

## INTRODUCTION

The subgranular zone (SGZ) of the hippocampal dentate gyrus (DG) is one of only a few regions of the mammalian brain that continues to exhibit neurogenesis into adulthood. Adult-born dentate granule cells (abDGCs) are continuously generated from a pool of largely quiescent neural stem cells that undergo proliferation, differentiation, and fate specification before maturing into neurons that are indistinguishable from developmentally derived dentate granule cells (DGCs) (Brunner et al., 2014; Espósito et al., 2005; Gu et al., 2012; Marin-Burgin et al., 2012; Temprana et al., 2015). These stem cells and their immature progeny are sensitive to environmental stimuli; their proliferation, development, and survival are regulated by multiple intrinsic and extrinsic factors, including experience, stress, and inflammation (Gonçalves et al., 2016a; Kempermann et al., 2015; Monje et al., 2003; Snyder et al., 2009; Vivar et al.). Numerous studies demonstrate that abDGCs are critical for maintaining the physiological activity of mature DGCs and contribute to hippocampus-dependent behaviors (Akers et al., 2014; Clelland et al., 2009; Clemenson et al., 2015; Deng et al., 2009, 2010, Ikrar et al., 2013, 2013; Ko et al., 2009; Lacefield et al., 2012; Nakashiba et al., 2012; Sahay et al., 2011a; Saxe et al., 2007; Singer et al., 2011; Tronel et al., 2012). While the specific role that immature DGCs play in hippocampal function, including the formation of memories, is not fully established, progress has been achieved through recent work focused on precisely modulating and measuring the activity of immature and mature DGCs within the DG of animals during behavior (Anacker et al., 2018; Danielson et al., 2016, 2017; GoodSmith et al., 2017; Hainmueller and Bartos, 2018; Hayashi et al., 2017; Kirschen et al., 2017; Leutgeb et al., 2007; Nakazawa, 2017; Pilz et al., 2016; Senzai and Buzsáki, 2017).

A key tool enabling many of these and other advances in *in vivo* neurophysiology is recombinant adeno-associated virus (rAAV). Wild-type AAV is a non-enveloped, single stranded DNA virus endemic to humans and primates and has been previously proposed to have no known pathogenicity. This replication-defective virus contains a 4.7 kb genome that includes the Rep and Cap genes and a pair of palindromic 145 bp inverted terminal repeats (ITRs). The Rep and Cap genes can be supplied in *trans* to create space for incorporating transgenes of interest, yielding the widely used rAAV, which retains only the ITRs from the original wild-type genome. In experimental neuroscience, rAAV is often used to deliver a variety of genetically encoded tools, including actuators and sensors of neuronal function, to specific cell types and brain regions. In addition, rAAV’s minimal viral genome and limited immunogenicity and toxicity have made it the vector of choice for human gene therapy (Büning and Schmidt, 2015; Choudhury et al., 2017; Hocquemiller et al., 2016; Hudry and Vandenberghe, 2019), including two FDA-approved therapies for disorders of the CNS (Hoy, 2019; Smalley, 2017).

Despite its safety profile, rAAV has increasingly been reported to demonstrate toxicity in some cell types (Bockstael et al., 2012; Hinderer et al., 2018; Hirsch et al., 2011; Hordeaux et al., 2018). However, the toxic effects of rAAV on abDGCs have previously not been assessed. Motivated by our own efforts to study the role of adult neurogenesis and the DG in learning and memory, we discovered that neural progenitor cells (NPCs) and immature neurons in the DG are highly susceptible to rAAV-induced death at a range of experimentally relevant titers (3 e11 gc/mL and above). This process appears to be cell autonomous and mediated by rAAV ITRs. Consistent with previous ablation studies, elimination of 4-week-old abDGCs by rAAV alters the activity of mature DGCs, resulting in DG hyperactivity (as indicated by cFOS expression). To circumvent this problem, we used the rAAV2-retro serotype (Tervo et al., 2016) to label DGCs in a retrograde manner, which avoids infection of susceptible cells and preserves adult neurogenesis. We demonstrate the utility of this delivery method by measuring the activity of mature DGCs *in vivo* using 2-photon calcium imaging.

## RESULTS

### rAAV eliminates abDGCs in a dose-dependent manner

We found that the delivery of calcium indicators using rAAV at doses equivalent to or below previously reported doses resulted in ablation of adult neurogenesis (**Fig. S1A**). This effect was robust regardless of vector production facility (Salk Institute Viral Vector Core, University of Pennsylvania Vector Core, Addgene), purification method (iodixanol, CsCl), capsid serotype (AAV1 & AAV8-shown; AAV9-not shown), promoter (CAG, Syn, CaMKIIa), and protein expression (GFP (**Fig. 1**), jRGECO1a and mCherry) at doses (>1 e12 gc/mL) typically required for the functional manipulation or visualization of DGCs *in vivo*.

**Figure 1.**
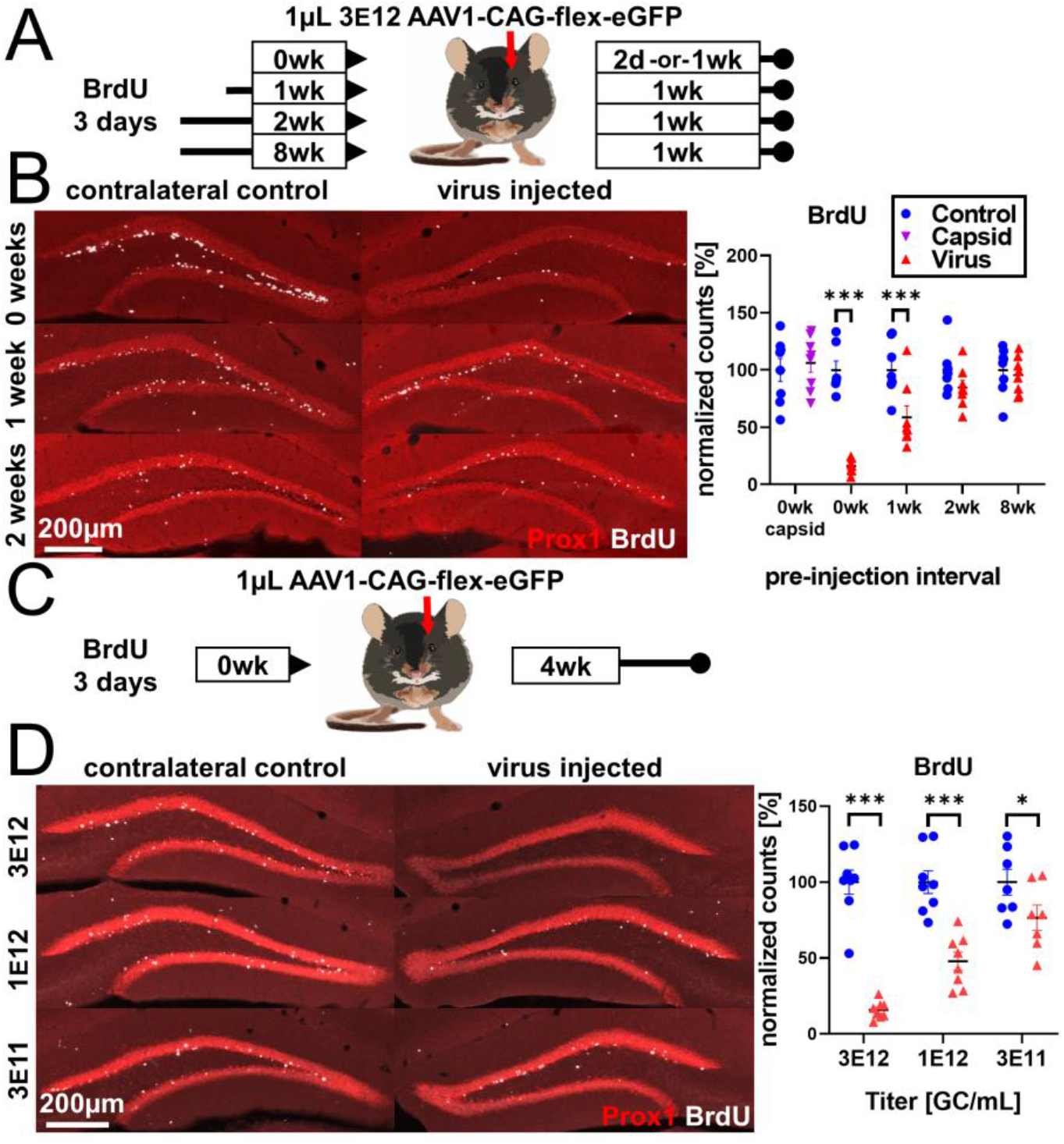
rAAV eliminates abDGCs in a dose-dependent manner. **(A)** Experimental design of rAAV injection into DG following indelible labeling of adult-born DGCs with BrdU. **(B, left)** Representative images showing Prox1 and BrdU used for quantification. **(B, right)** abDGCs birth-dated with BrdU 3 days immediately preceding viral injection show near complete elimination following rAAV injection; cells born 1 week before viral injection are reduced ~50%. Cells born 2 weeks or more, or injection of empty AAV viral capsid, demonstrate no reduction. **(C)** Experimental design of dose-dependent attenuation of abDGCs by rAAV. **(D, left)** Representative images showing Prox1 and BrdU used for quantification. **(D, right)** Immediately following birth-dating of BrdU+ abDGCs, a near complete ablation of BrdU+ cells is seen in the DG injected with 1μL 3 e12 GC/mL rAAV, partial ablation of BrdU+ cells results from the injection of 1μL 1 e12 GC/mL rAAV, and a small but significant reduction of adult neurogenesis results from injection of 1 μL 3 e11 GC/mL rAAV. This pattern is matched by Tbr2 expression (Figure S1B). All data are presented as mean ± s.e.m, significance reported as: * p < 0.05, *** p < 0.001.

To systematically quantify the effect of rAAV transduction on abDGCs, we chose to inject a widely available, minimally expressing cre-recombinase-dependent virus (AAV1-CAG-flex-eGFP, U. Penn. & Addgene #51502) in non-cre-expressing wild type C57BL/6J mice to mitigate any contributions from toxicity that might be attributed to protein expression. Mice received daily intraperitoneal injections of 5-bromo-2’-deoxyuridine (BrdU) for 3 days to label dividing cells; then 1μL of 3e12 gc/mL rAAV was injected unilaterally into the DG immediately (0d), or 1 or 2 weeks later (schematic in **Fig. 1A**). Cell survival on the virus-treated side depended on the age of the cells when virus was delivered (F_treatment x time_(3,27)=29.0, p<0.001; **Fig. 1B** last 4 columns). Cells that were 2 days old and younger at the time of injection were almost completely eliminated within 48 hours (−83.9% ± 6.7%, p <0.001, all textual results reported as change relative to non-injected contralateral DG +/− standard error of the mean difference, unless stated otherwise). Cells that were 7-9 days old were partially protected (−41.3% ± 6.3%, p <0.001), whereas cells that were 14-16 days old were largely protected with variable but non-significant loss (−15.4% ± 6.3%, n.s.). Mature abDGCs approximately 8 weeks old also did not demonstrate significant loss (4.5% ± 6.3%, n.s.; **Fig. 1B** last column). In the same tissue, Tbr2+ intermediate progenitors were lost even when mature BrdU+ cells were spared, demonstrating that the virus did not lack toxicity in these animals (F_treatment x time_(3,27)=3.0, p < 0.05; 0 weeks: −80.2% ± 9.1%, p<0.001; 1 weeks: −76.9% ± 8.5%, p<0.001; 2 weeks: −82.6% ± 8.5%, p<0.001; 8 weeks: −50.8% ± 8.5%, p<0.001; **Fig. S1B**). To determine the effect of viral attachment and penetration in rAAV-induced toxicity, we injected 1μL of high titer (3.7 e13 capsids/mL) empty AAV viral capsid into the DG (**Fig. 1B** first column, **Fig S1C**). At 1week post-injection, there was no effect on BrdU+ cells (2 days old at the time of empty capsid injection) relative to the contralateral control DG (F_treatment x time_(1,14)=6.5, p<0.05; 1 week post-injection: 6.2% ± 3.2%, n.s., **Fig. 1B** first column; although there was a mild decrease 4 weeks post-injection: −14.5% ± 7.4%, p<0.05, which did not recapitulate the severe loss observed with intact virus; **Fig S1C**).

We then assessed the effect of titer on rAAV-induced cell loss. We labeled abDGCs for 3 days with BrdU and injected 1μL of either 3 e12 gc/mL, 1 e12 gc/mL, or 3 e11 gc/mL rAAV on the final day of BrdU labeling (schematic in **Fig. 1C**). Cell loss increased with increasing titer of virus injected (F_treatment x titer_(2,20)=19.2, p<0.001). A nearly complete ablation of BrdU+ cells was seen in the DG injected with 3 e12 gc/mL rAAV (−84.3% ± 6.7%, p <0.001), whereas partial ablation of BrdU+ cells resulted from the injection of 1 e12 gc/mL rAAV (−52.1% ± 6.7%, p <0.001), and a small but significant reduction of adult neurogenesis resulted from injection of 3e11 gc/mL rAAV (−23.4% ± 7.2%, p <0.05; **Fig 1D**). This dose-dependent pattern was matched by reductions in immature neuron marker DCX expression (**Fig. S1D**).

### Developmental stage determines susceptibility to rAAV-induced cell loss

After determining the differential response of abDGCs to rAAV based on post-mitotic age, we determined which population of NPCs was susceptible to rAAV-induced loss. To accomplish this, we varied the post-injection interval and measured canonical early (Sox2), middle (Tbr2), and late (DCX) histological markers associated with abDGC development. Mice were unilaterally injected with 1μL of 3 e12 gc/mL rAAV and sacrificed at 2 days, 1 week, or 4 weeks post-injection (schematic in **Fig. 2A**). The number of Sox2+ cells within the SGZ was modestly decreased (F_treatment_(1,19)=15.5, p<0.001; F_time_(2,19)=1.6, n.s.; F_treatment x time_(2,19)=2.7, n.s.; **Fig. 2B,C**). In contrast, Tbr2+ intermediate progenitor cells were almost entirely lost and did not show signs of recovery by 4 weeks post-injection (F_treatment_(1,19)=129.2, p<0.001; F_time_(2,19)=0.1, n.s.; F_treatment x time_(2,19)=0.2, n.s.; **Fig. 2B, D**). Injection of empty AAV viral capsid (**Fig. 2E)** did not result in a similar reduction of Tbr2+ cells at 1 or 4 weeks post-injection (F_treatment_(1,10)=6.4E-5, n.s. F_time_(1,10)=0.3, n.s.; F_treatment x time_(1,10)=1.8, n.s; **Fig. 2F**). Expression of the late premitotic and immature neuronal marker DCX showed progressive decline until near complete loss at 4 weeks post-injection (F_treatment x time_(3,27)=12.8, p<0.001; 2 days: −27.7% ± 9.0%, p<0.01; 1 week: −58.7% ± 8.4%, p<0.001; 4 weeks: −92.0% ± 9.0%, p<0.001; **Fig. 2B, G**) and did not show signs of recovery 3 months post-injection (−68.7% ± 8.0%, p<0.001; **Fig. 2G**, last column). DCX expression was not altered following injection of empty AAV viral capsid (F_treatment_(1,14)=3.9, n.s.; F_time_(1,14)=0.7, n.s.; F_treatment x time_(1,14)=1.3, n.s.; **Fig S2A**).

**Figure 2.**
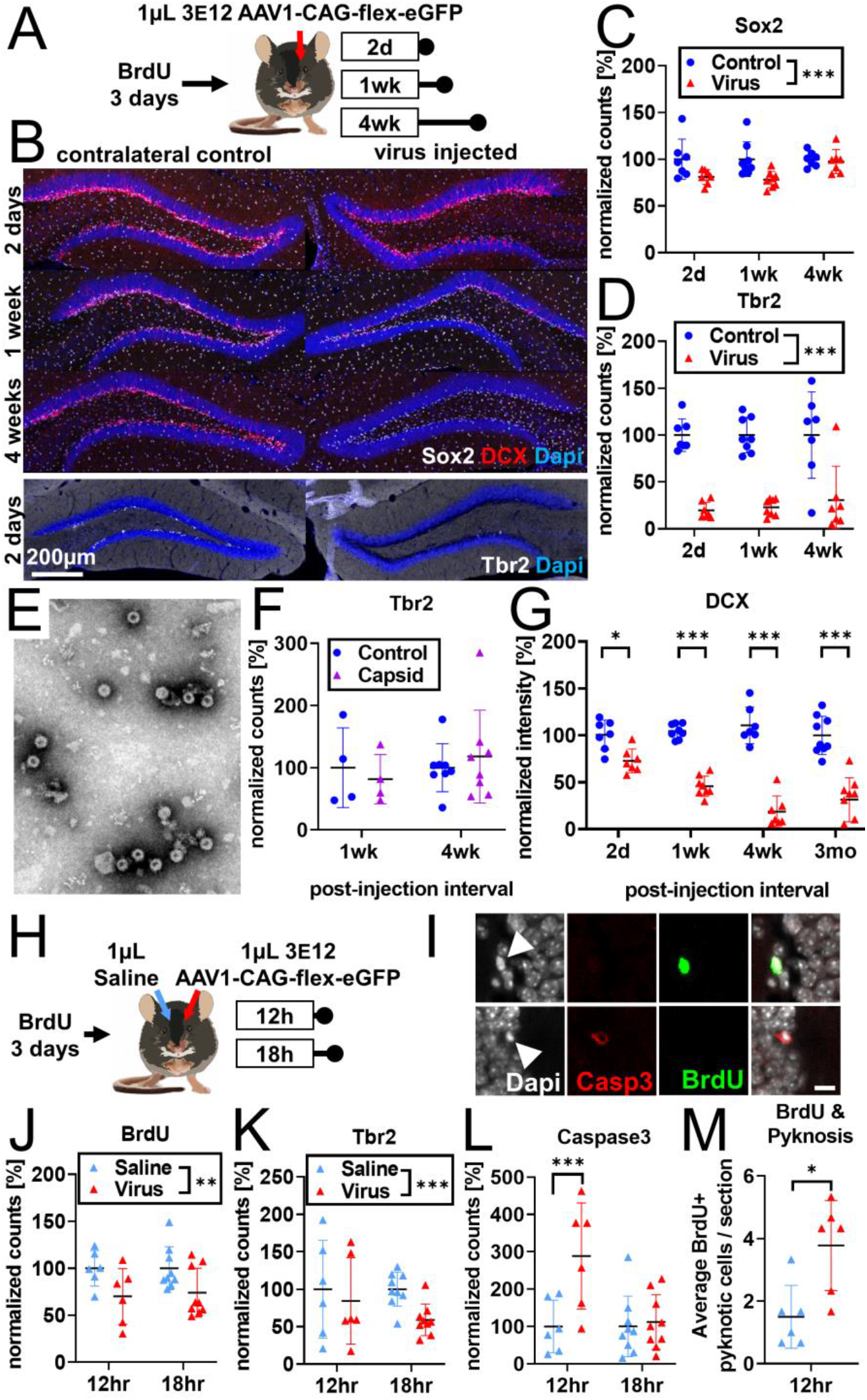
Developmental stage determines susceptibility to rAAV-induced cell loss. **(A)** Experimental design to assess the effect of rAAV post-injection interval on the survival of different NPC types. Following labeling with BrdU, mice are injected unilaterally with 1 μL of 3 e12 GC/mL rAAV and sacrificed at 2 days, 1 week, and 4 weeks. **(B)** Representative histological staining of progenitor and immature neuronal markers Sox2 (white, upper panels), DCX (red, upper panels), and Tbr2 (white, lower panels) following rAAV injection. **(C)** Sox2+ neural stem cell numbers within the SGZ are reduced by ~20% 2 days and 1 week following rAAV injection but not at 4 weeks post-injection. **(D)** The majority of immediate early progenitor Tbr2+ cells are lost within 2 days of rAAV injection and do not recover by 4 weeks post-injection. **(E)** Representative image of empty AAV viral particles (“empty capsid”) from cryo-electron microscopy used to quantify the number of viral particles. **(F)** Tbr2+ intermediate progenitors are preserved following injection of empty viral capsid. **(G)** Immature neuronal marker DCX intensity shows progressive decline until complete loss at 4 weeks post-injection. DCX intensity shows no recovery at 3 months post-injection. **(H)** Experimental design for acute time-line of rAAV-induced cell loss. Following labeling with BrdU, 1 μL 3 e12 GC/mL rAAV and saline control are injected into opposite sides of the DG; mice are sacrificed at 12 and 18 hours. **(I)** DNA condensation and nuclear fragmentation (pyknosis and karyorrhexsis, white arrowheads) are assessed with BrdU (green), Caspase 3 activation (red), and DAPI. **(J)** BrdU+ cells show variable decline 12 hours after rAAV injection and significant decline at 18 hours relative to saline control. **(K)** Tbr2+ intermediate progenitors show significant decline by 18 hours following rAAV injection. **(L)** Caspase-3+ apoptotic cells were increased relative to saline-injected controls at 12 hours. **(M)** BrdU+ cells exhibit a significant increase in pyknosis 12 hours after rAAV injection. Increase in Caspase 3 activation of pyknotic cells is seen at 12 hours. All data are presented as mean ± s.e.m, significance reported as: * p < 0.05, ** p < 0.01, *** p < 0.001.

These findings suggest that, despite an initial sensitivity of some Sox2+ cells to rAAV transduction, this largely quiescent neural progenitor pool remains mostly intact. Instead, a rapid loss of proliferating Tbr2+ intermediate progenitors by 2 days drives much of the rAAV-induced toxicity, including the progressive loss of the DCX+ population that is observed as these cells differentiate into DCX-neurons and decline in number over time. To test the effect of increasing the size of the proliferating NPC population on rAAV-induced toxicity, we housed mice in an enriched environment with running wheels (van Praag et al., 1999). Nearly all of the BrdU+ dividing cells gained from enrichment and exercise were eliminated by rAAV (F_housing_(1,14)=8.7, p<0.01; F_treatment_(1,14)=124.0, p<0.001; F_housing x treatment_ (1,14)=9.4, p<0.01; home cage: −79.5% ± 13.9%, p<0.001; enriched environment: −139.9% ± 13.9%, p<0.001; **Fig. S2B**). Tbr2+ cells were unaffected by enrichment and eliminated by rAAV regardless of housing type (F_housing_(1,14)=0.8, n.s.; F_treatment_(1,14)=147.1, p<0.001; F_housing x treatment_(1,14)=0.5, n.s.; **Fig. S2C**). These results indicate that proliferating cells are the primary target of the virus.

Given the extensive and rapid loss of BrdU+ (**Fig. 1B,C**) and Tbr2+ (**Fig. 2B,D**) cells, we designed an acute time-course experiment to determine the mechanism of rAAV-induced cell loss (schematic **Fig. 2H,I**). Following labeling with BrdU, animals were injected with 1μL of 3e12 rAAV into one dorsal DG and 1μL saline into the contralateral DG to control for the acute effect of surgery- and injection-induced inflammation and tissue damage. rAAV-injected DGs had a modest decrease in BrdU+ cells at 12 and 18 hours relative to their contralateral saline-injected control (F_treatment_(1,13)=13.9, p<0.01; F_time_(1,13)=0.04, n.s.; F_treatment x time_(1,13)=0.07, n.s.; **Fig. 2J**). The same decrease was seen in Tbr2+ cells (F_treatment_(1,13)=16.5, p<0.001.; F_time_(1,13)=0.4, n.s.; F_treatment x time_(1,13)=3.3, n.s.; **Fig. 2K**). Cell loss was accompanied by an increased number of Caspase-3+ apoptotic cells relative to saline-injected controls at 12 hours (F_treatment x time_(1,13)=21.2, p<0.001; 12h treatment: +188.6% ± 29.8%, p <0.001; 18h treatment: 11.7 ± 24.3%, n.s.; **Fig. 2L**). Therefore, we determined that 12 hours would be a suitable time point to investigate nuclear changes in dying cells, before extensive cell loss had occurred. Condensed and fragmented chromatin (pyknosis and karyorrhexis) was identified in conjunction with BrdU and Caspase-3 (**Fig. 2I**). Increased numbers of pyknotic and karyorrhexic cells were identified in rAAV-injected DGs relative to their saline-injected contralateral controls (14.7 ± 6.3 additional cells/section, p<0.05, **Fig. S2D**). A significant increase in pyknosis was also seen in BrdU+ proliferating cells (2.3 ± 0.7 additional cells/section, p<0.05; **Fig. 2M**). Pyknotic cells were more likely to be Caspase 3+ following rAAV injection relative to saline controls (7.7 ± 1.4 additional cells/section, p<0.01; **Fig. S2E**); though BrdU+ Caspase-3+ pyknotic cells were particularly rare (n=4 of 887 cells, all in rAAV-injected DG). Taken together, these findings suggest that rAAV increases programmed cell death of dividing and recently divided cells in the DG.

Both systemic and local inflammation are known to negatively impact adult neurogenesis (Ekdahl et al., 2003; Monje et al., 2003). Therefore, we investigated whether rAAV-induced cell loss could be explained by inflammation resulting from rAAV infection. In contrast to the rapid loss of NPCs (**Fig. 2**), expression of the microglial marker, Iba1 was not increased until 4 weeks post-injection (F_treatment x time_(2,19)=54.6, p<0.001; 2 days:18.6% ± 10.3%, n.s.; 1 week: 9.0% ± 9.7%, n.s.; 4 weeks: 132.4% ± 10.34%, p<0.001; **Fig. S2 F,H**), and no obvious change in microglial morphology was observed at 2 days or 1 week relative to contralateral control. At 4 weeks, microglia exhibited an amoeboid morphology, indicative of active inflammation. Similarly, expression of the astrocyte marker GFAP was unchanged in the SGZ and hilus at 2 days post-injection, slightly increased at 1 week, and greatly increased at 4 weeks (F_treatment x time_(2,19)=91.0, p<0.001; 2 days: 21.6% ± 8.7%, n.s.; 1 week:25.2% ± 8.1%, p<0.05; 4 weeks: 165.5% ± 8.7%, p<0.001; **Fig. S2G,H**). To further rule out a bystander effect, we injected 30 nL of 5 e12 gc/mL AAV1-Syn-NES-jRGECO1a into the DG. Histology demonstrated incomplete loss of DCX labeling that faithfully followed the boundaries of transgene expression, where uninfected cells located microns away from those expressing jRGECO1a were spared **(Fig. S2I**). These findings suggest that AAV-induced toxicity may be cell autonomous and is unlikely to be mediated by astrocyte- or microglia-activated immune responses or by inflammatory signals and other changes within the niche (Ekdahl et al., 2003). Additionally, we found that rAAV-induced toxicity was not altered in Sting knockout mice and, therefore, likely not dependent on foreign nucleic acid detection through Sting-mediated pathways (BrdU: −90.2% ± 14.2, p<0.001; Tbr2: −88.7% ± 12.9%, p<0.001; **Fig. S2J**).

### rAAV induces toxicity in NPCs *in vitro*

To further explore whether rAAV-mediated toxicity is cell-autonomous, we developed an *in vitro* assay to study rAAV-induced elimination of NPCs. Mouse NPCs were plated, administered rAAV with a multiplicity of infection (MOI) of 1 e4 TO 1 e7 or water control, and chronically imaged to examine cell survival and proliferation. Dose-dependent inhibition of NPC proliferation and cell death was most profound in NPCs infected with rAAV 1 e7 MOI and moderate in NPCs infected with 1 e6 MOI. Within 24 hours of rAAV application, NPCs infected with 1 e7 MOI had ceased to proliferate, whereas application of 1 e6 MOI resulted in comparatively slower proliferation compared to H20 control. Infections with 1 e5 and 1 e4 MOI were nearly indistinguishable from H2O control (F_treatment_(4,665)=1039, p<0.001; F_time_(18,665)=1341, p<0.001; F_treatment x time_(72,665)=52.7, p<0.001**; Fig. 3A,C**). Cell death, visualized by permeability to propidium iodide, also showed a dose-dependent increase. NPCs infected with 1 e7 MOI showed the most significant increase in cell death, whereas NPCs infected with 1 e6 MOI showed an intermediate increase. NPCs infected at 1 e5 and 1 e4 MOI were indistinguishable from H2O control (F_treatment_(4,665)=9256, p<0.001; F_time_(18,665)=1127, p<0.001; F_treatment x time_(72,665)=323.5, p<0.001**; Fig. 3B,C**).

**Figure 3.**
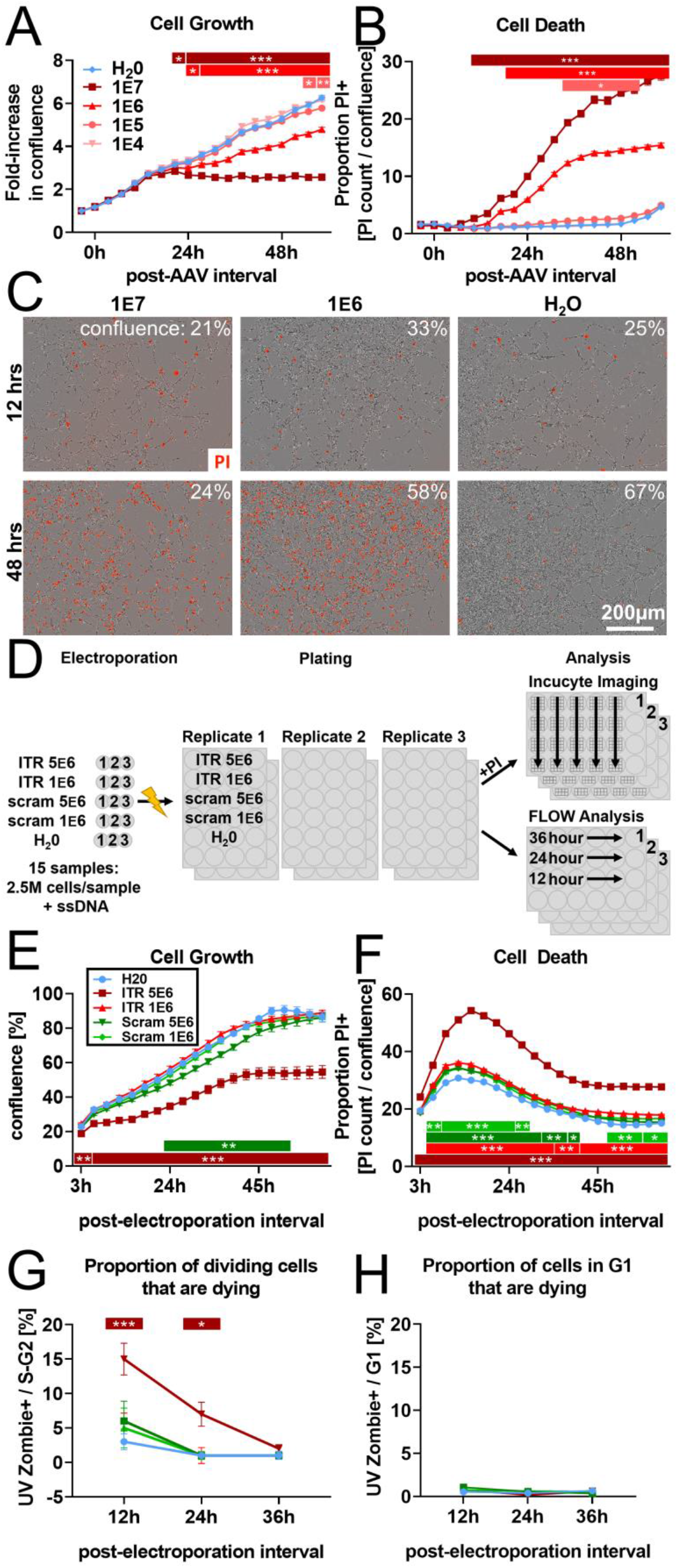
rAAV induces toxicity in NPCs *in vitro*. **(A)** Dose-dependent inhibition of NPC proliferation by rAAV; initial multiplicity of infectivity (MOI) of 1 e7 viral particles/cell arrests mNPC proliferation by 24 hours. MOI of 1 e6 results in slower proliferation relative to H2O control, MOI of 1 e5 and lower are indistinguishable from H2O control. **(B)** Dose-dependent rAAV-induced death; MOI of 1 e7 and 1 e6 result in increased proportion of propidium iodine+ NPCs. **(C)** Representative images showing confluence (brightfield) and propidium iodide penetration (red) into NPCs 12 and 48 hours post-viral transduction for MOI of 10^7^, 10^6^, and for H20 control. **(D)** Experimental design for 145bp ssDNA AAV ITR electroporation. Mouse NPCs are electroporated with 5 e6 or 1 e6 copies of 145bp ssDNA AAV ITR or scrambled ITR sequence control per cell and plated for FLOW analysis time course or treated with propidium iodide for imaging time course on Incucyte S3. Statistical significance shown as post-hoc Tukey’s multiple comparison test relative to H20 (also see Table S1). **(E)** Electroporation of 5 e6 ITR is sufficient to result in cell loss within hours of electroporation and arrest of proliferation by 40 hours. 5 e6 scrambled ITR shows slight decrease in confluence relative to 1 e6 scrambled ITR; doses of 1 e6 ITR and H2O control are indistinguishable from 1 e6 scrambled ITR. **(F)** Electroporation of ITR is sufficient to induce greater levels of cell death at higher concentration of 5e6 copies/cell. **(G)** FLOW analysis demonstrates dose-dependent effect of rAAV ITR on replicating NPCs whereby cells electroporated with 5 e6 ITR in S- and G2-phase are dying and permeable to UVZombie at 12 hours post-electroporation. **(H)** NPCs in G1 represent the vast majority of cells (Figure S3I) and are not substantially dying or permeable_to UVZombie, regardless of treatment. All data are presented as mean ± s.e.m, significance reported as: * p < 0.05, ** p < 0.01, *** p < 0.001.

We then examined whether the minimum components of the AAV genome required for viral encapsulation, the 145bp ITRs, were sufficient to induce cell death as previously reported in embryonic stem cells (Hirsch et al., 2011). NPCs were electroporated with “high” (5 e6 copies/cell) and “low” (1 e6 copies/cell) doses of 145bp rAAV ITR ssDNA, scrambled ITR sequence control, or water and plated for imaging (as above) or for FACS analysis (schematic in **Fig. 3D**). In the high-dose ITR condition, NPCs were significantly decreased by 6 hours post-electroporation (F_treatment_(4,200)=630.2, p<0.001; F_time_(19,200)=635.0, p<0.001; F_treatment x time_(76,200)=6.0, p<0.001; 6 hour ITR 5 e6 vs H20: −8.1% ± 2.5, p<0.05) and had ceased expansion by 40 hours (**Fig. 3E**). Low-dose ITR and low-dose scramble groups were indistinguishable from H20. Although a transient decrease in the high-dose scrambled condition relative to H20 was observed, this decrease was minimal compared to the effect of high-dose ITR (**Fig. 3E** and **Table S1**). The number of dying propidium iodide+ cells increased in all groups in the first 24 hours following electroporation (F_treatment_(4,200)=3729, p<0.001; F_time_(19,200)=1219, p<0.001; F_treatment x time_(76,200)=19.3, p<0.001**; Fig. 3F**), with the proportion of dying cells decreasing as confluence increased during the experiment. This proportion was substantially greater in the high ITR condition relative to H20 (**Fig. 3F** and **Table S1**). Both low- and high-dose scrambled groups had a slight increase in cell death relative to H20 that was minimal compared to the effect of high-dose ITR. FACS analysis at 12, 24, and 36 hours showed the proportion of cells in S/G2 phase that were dying (UVZombie+) was greatly increased at 12 and 24 hours in the high ITR condition but not in the other experimental groups relative to H20 control (F_treatment_(4,30)=9.5, p<0.001; F_time_(2,30)=23.4, p<0.001; F_treatment x time_(8,30)=2.2, p<0.05; 12h ITR 5e6 vs H20 +12.0% ± 2.0%, p <0.001, **Fig. 3G**). The proportion of non-replicating cells that were dying was <1% in all groups (Fig. 3H).

### AAV retro serotype permits studies of DGC activity *in vivo* without ablating adult neurogenesis

To determine the functional consequence of AAV-induced ablation of neurogenesis on DG activity, animals were injected with 1μL of 3 e12 gc/mL, 1 e12 gc/mL, or 3 e11 gc/mL rAAV and exposed 4 weeks later to a novel environment (NE) prior to sacrifice. DGC expression of the immediate early gene cFOS was used to quantify the effect of AAV on DG activity. Consistent with previous studies examining the effects of manipulating adult neurogenesis on DG activity (Ikrar et al., 2013), mature DGC cFOS activation showed an inverse relationship with the level of adult neurogenesis (F_treatment x time_(2,20)= 11.4, p<0.001; **Fig. 4A,B**). Injection with 1μL 3 e12 gc/mL rAAV, which ablates over 80% of BrdU+ cells (**Fig. 1D**), resulted in the largest increase in mature DGC cFOS activation (81.6±13.6 additional cells per section, p<0.001); 1μL 1 e12 gc/mL rAAV injection resulted in a moderate and more variable increase (40.0 ± 13.6 additional cells per section, p<0.05), and 1μL 3 e11 gc/mL rAAV injection demonstrated no significant change on average (13.3 ± 14.5 additional cells per section, n.s.). cFOS activation and loss of BrdU+ cells were significantly correlated (slope =0.26, R^2^=0.59, p<0.001; **Fig. S3A**). These results demonstrate that, while viral titers producing severe ablation of neurogenesis have the most severe effect on DG activity, titers that do not completely eliminate neurogenesis still have an impact on the network.

**Figure 4.**
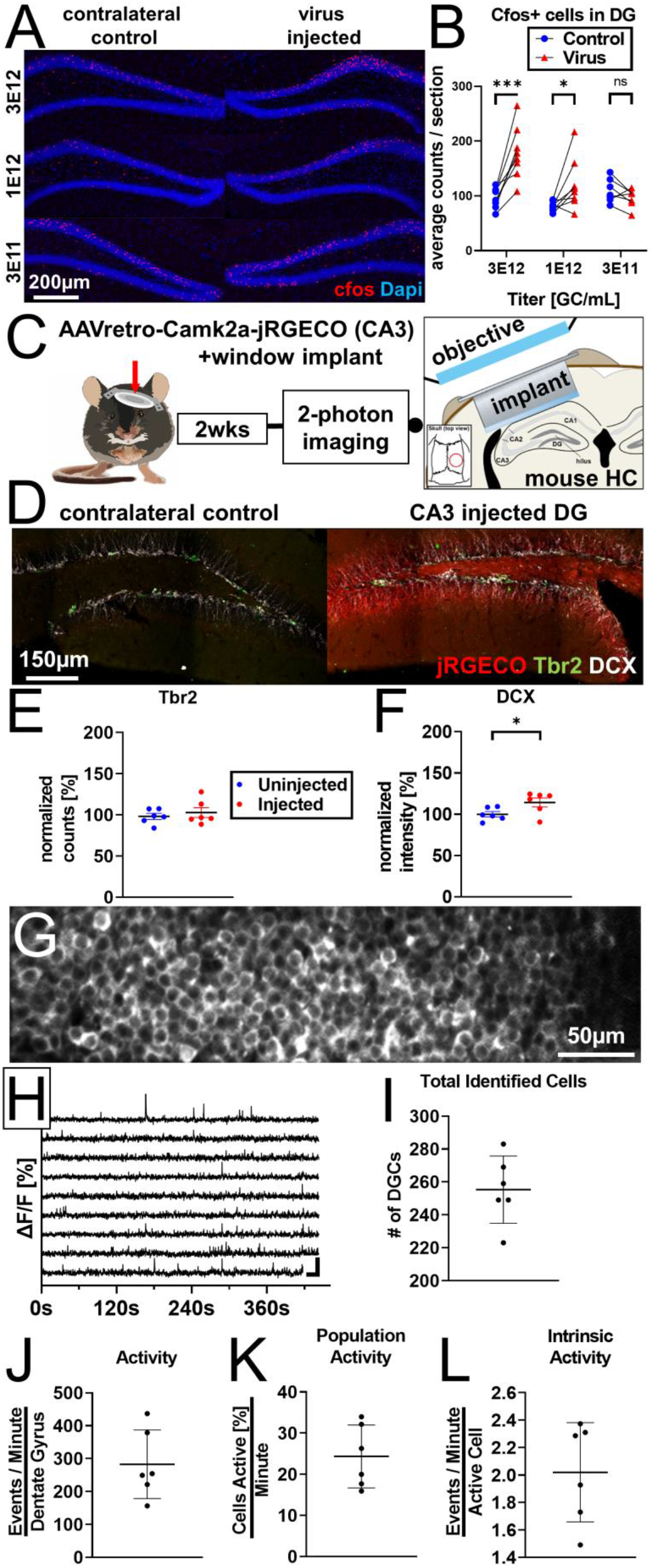
AAV retro serotype permits studies of DGC activity *in vivo* without ablating adult neurogenesis. **(A)** Representative images showing cFos and Dapi used for quantification. **(B)** Mature DGCs are hyperactive following rAAV-induced cell loss in a dose-dependent manner for injected titers between 3e12 and 3e11. abDGC knockdown efficiency is significantly correlated with cFOS activation in mature DGCs (Figure S3A). **(C)** Experimental design for 2-photon imaging of DG utilizing AAV retro. 800 nL of 3 e12 GC/mL AAVretro-CaMKIIa-jRGECO1a is injected into CA3, a cranial window is implanted, and mice undergo 2-photon calcium imaging 2 weeks later. **(D)** Representative images show Tbr2+ and DCX+ cells are intact in animals injected with AAV retro in CA3. **(E)** Quantification of Tbr2 intermediate progenitors demonstrating adult neurogenesis is intact in AAV retro-injected animals. **(F)** Quantification of DCX staining demonstrates adult neurogenesis is intact in AAV retro-injected animals. **(G)** Representative field of view maximum projection for 2-photon calcium imaging showing cytoplasmic expression of jRGECO1a in >200 DGCs within a field of view. **(H)** Representative calcium traces of 10 randomly selected neurons from the same animal shown above. **(I)** Total number of identified DGCs in each DG is similar across animals. **(J-L)** Total activity of these cells was demonstrated to be sparse, with approximately one quarter of all cells active in any given minute while imaging, and only a few calcium transients per active cell a minute. Data are presented as mean ± s.e.m when comparing between groups in C-D and mean ± s.d. when describing variability within groups in G-J, significance reported as: * p < 0.05, ** p < 0.01, *** p < 0.001.

Given the importance of adult neurogenesis in regulating population activity in the DG, we sought a method that would permit 2-photon calcium imaging of *in vivo* network activity within the DG without ablating neurogenesis. AAV retro is a designer AAV variant capsid optimized to be taken up by axonal projections (Tervo et al., 2016). We used AAV retro (AAVretro-CaMKIIa::NES-jRGECO1a) to deliver the red calcium indicator jRGECO1a to DGCs in a retrograde fashion by injecting virus into the dorsal CA3, where their axons (“mossy fibers”) terminate. This delivery method allowed us to avoid infecting immature DGCs whose mossy fiber projections do not reach CA3 until after 2 weeks of age, a time point when these neurons also demonstrate decreased susceptibility to rAAV-induced toxicity (**Fig. 1B**). Following viral injection, a cranial window was implanted to permit visualization of DGC activity via 2-photon calcium imaging (schematic in **Fig. 4C**).

Histology performed following 2-photon calcium imaging confirmed that adult neurogenesis was intact 2 weeks after injection relative to the uninjected contralateral hippocampus (Tbr2: +4.7% ± 7.0%, t(5)=0.7, n.s. **Fig. 4D,E**; DCX +14.3 ± 6.3%, t(5)=4.3, p<0.01, **Fig. 4D,F**). Spontaneous calcium activity was recorded in DGCs and quantified in these mice under head fixation on a cylindrical treadmill in the dark (**Fig. 4G,H**). A similar number of DGCs was identified across animals (255.3 ± 20.5 cells/DG; **Fig. 4 I**). Total activity of these cells was demonstrated to be sparse (282.9 ±104.5 events/minute; **Fig. 4J**). Approximately one quarter of the cells were active in any given minute during imaging (24.3% ± 7.6% cells active/minute; **Fig 4K**), with a small number of calcium transients per active cell observed per minute (2.0 ± 0.4 events/active cell/minute; **Fig. 4L**). The limited activity of DGCs measured in our study with AAV retro is consistent with previous imaging studies in the DG (Danielson et al., 2016; Pilz et al., 2016). Due to the extensive inflammation that arises 4 weeks after viral injection (**Fig. S1A, Fig S2G-I**), we were unable to image the DG at 4-weeks post-viral injection, when abDGCs are highly active and known to contribute to DG activity (**Fig. 4A**). Despite this limitation, a small suppressive effect of AAV1 injection on DG activity was already evident by 2-photon imaging 2-weeks post injection (**Fig. S3**), when DG imaging is often initiated (Danielson et al., 2016; Hainmueller and Bartos, 2018; Pilz et al., 2016). No change in cFOS expression due to AAV1 was observed at this earlier time point (**Fig. S3K**). Collectively, these findings demonstrate that 1) delivery of AAV directly into the DG ablates neurogenesis and influences activity in this network, particularly at time points when abDGCs make a strong contribution to DG function, and 2) AAV retro enables delivery of transgenes to the DG, permitting studies of DGC function *in vivo* while leaving adult neurogenesis intact.

## DISCUSSION

### A developmental window for sensitivity to rAAV-induced toxicity

We demonstrate that adult murine NPCs and immature neurons up to approximately 1 week of age are eliminated by rAAV in a dose-dependent fashion (**Fig. 1**). The doses demonstrated to ablate neurogenesis are within or below the range of experimentally relevant titers commonly injected into the mouse DG, 1.5 e12 to 3.6 e13 gc/mL (Anacker et al., 2018; Castle et al., 2018; Danielson et al., 2016, 2017; Gong and Zhou, 2018; Hashimotodani et al., 2017; Hayashi et al., 2017; Kaspar et al., 2002; Kirschen et al., 2017; Liu et al., 2012; McAvoy et al., 2016; Ni et al., 2019; Pilz et al., 2016; Ramirez et al., 2013; Raza et al., 2017; Redondo et al., 2014; Senzai and Buzsáki, 2017; Swiech et al., 2015; Zetsche et al., 2017). This rAAV-induced cell death is rapid and persistent; BrdU-labeled cells and Tbr2+ intermediate progenitors begin to die within 12 to 18 hours post-injection and are eliminated by 48 hours (**Fig. 2**). Under physiological conditions, the Tbr2+ population replenishes DCX+ progenitors and immature neurons, which can retain expression of the DCX protein in the mouse for more than 3 weeks post mitosis (Kempermann et al., 2003). Upon administration of rAAV, many of the DCX+ neurons are postmitotic and initially spared but are not replenished by the ablated Tbr2+ population, explaining the delayed and progressive loss of the DCX+ pool in response to rAAV infection. Neither of these populations showed evidence of recovery when assessed several months post-injection (**Fig. 1B, 2G**). Interestingly, the total number of Sox2+ cells, composed of Type I and IIa NPCs, was relatively unaffected by rAAV, consistent with this population being largely quiescent (**Fig. 2 B,C**). The fact that Sox2+ cells were mostly preserved in our experiments might explain why Type I cells are visualized in studies utilizing a large range of AAV titers, including titers greater than those reported here (Crowther et al., 2018; Kotterman et al., 2015; Ojala et al., 2018; Pulicherla et al., 2011; Song et al., 2012).

*In vitro* application of rAAV or electroporation of AAV2 ITRs is sufficient to induce arrest of proliferation and cell death, pointing toward a cell autonomous process (**Fig. 3**). This is consistent with our findings that the timing of rAAV-induced apoptosis and cell loss (hours to days) is not commensurate with the time course or spatial extent of inflammation (**Fig. S2G-I**) and is independent of exposure to empty capsids (**Fig. 1B, S1A, 2F, S2A**). Analysis with FACS demonstrated that administration of high-dose ITR oligonucleotides resulted in a disproportionate loss of dividing cells (**Fig. 3G,H**). To a large extent, this seems to recapitulate *in vivo* experiments, which demonstrate that a developmental window exists where actively dividing Type II NPCs and recently post-mitotic immature neurons are sensitive to rAAV-induced cell death, whereas rarely dividing neural stem cells and mature neurons flanking this window are significantly less affected. Note that the persistence of Sox2 expression in dividing Type 2a progenitors may account for the initial modest decrease seen in the number of Sox2+ cells (Gonçalves et al., 2016b; Kempermann et al., 2015). Why these cells do not replenish the Tbr2+ pool following rAAV infection remains unknown. One possibility is that Type I cells infected with rAAV undergo delayed apoptosis upon entering the cell cycle, precluding the recovery of neurogenesis after infection.

### rAAV as a model system for viral toxicity in the developing CNS

Infections involving a number of viruses, including cytomegalovirus (CMV), rubella, varicella-zoster, herpes simplex, human immunodeficiency virus (HIV), and Zika, have been implicated in the pathogeneses of microcephaly, the abnormal development of the cerebral cortex resulting in small head size. There is evidence that these viruses cause microcephaly through the elimination of NPC populations (Devakumar et al., 2018) in HIV (Balinang et al., 2017; Evans et al., 2016; Lawrence et al., 2004; Rothenaigner et al., 2007; Schwartz et al., 2007), CMV (Luo et al., 2010; Rolland et al., 2016; Teissier et al., 2014), and Zika virus (Garcez et al., 2016; Qian et al., 2016; Wen et al., 2017) among others. However, the complex biology of these viruses and their neural progenitor targets has precluded elucidation of a precise mechanism. Perhaps the best studied among these is Zika virus, which, similar to rAAV, attenuates proliferation and neurogenesis in the adult mouse DG, demonstrating a marked loss of EdU+ cells following infection (Li et al., 2016). These findings resemble the loss of proliferating cells in brain organoids and other models of the developing nervous system in response to Zika infection (Cugola et al., 2016; Dang et al., 2016; Garcez et al., 2016; Qian et al., 2016). Collectively, these studies have focused on Zika virus’ selective tropism for Sox2+ and Nestin+ NPCs over Tbr2^+^ and other immature cell types in the brain. However, these studies typically measure the fraction of Zika-infected cells that expresses Sox2, Tbr2, and other immature markers but do not track changes in the total size of these populations resulting from Zika infection. These measurements could have missed a rapid elimination of Tbr2 cells, which in our measurements was apparent within 48 hours of rAAV infection (**Fig 2D,K**). Alternatively, the proliferative capacity and thus the susceptibility of the Sox2+ population to viral infection could differ between developmental models and the adult DG, where in adult neurogenesis Sox2+ cells are largely quiescent and perhaps less sensitive to viral-induced cell death (Wu et al., 2018). Further studies are needed to discern the downstream events that lead to viral toxicity and whether this heterogeneous collection of viruses kills dividing NPCs through a common pathway. rAAV, with its exceedingly simple genome, broad tropism, and inability to replicate, offers a tractable model system to dissect the molecular events underlying this important phenomenon.

### Implications for DG and hippocampal function

Both theoretical and experimental studies indicate that the DG is involved in hippocampus-dependent behavioral pattern separation and pattern completion (Chawla et al., 2005; Deng et al., 2013; Lacy et al., 2011; Leutgeb et al., 2007; Marr, 1971; McClelland et al., 1995; Treves and Rolls, 1994; Yassa and Stark, 2011). More recent studies implementing genetically encoded tools, often delivered via rAAV, provide striking evidence for the role of memory engram representations in behavioral pattern separation and completion in the DG (Bernier et al., 2017; Danielson et al., 2016; Liu et al., 2012; Ramirez et al., 2013; Redondo et al., 2014) and highlight the role of this circuit in affective disorders and stress responses (Anacker et al., 2018; Li et al., 2017; Ni et al., 2019; Ramirez et al., 2015; Shuto et al., 2018; Wang et al., 2017). Moreover, DG activity and computations appear to depend on the addition of abDGCs (Clelland et al., 2009; Ikrar et al., 2013; Sahay et al., 2011a), which attenuate the activity of mature DGCs. This net inhibition on mature DGCs is thought to enhance pattern separation by selectively suppressing competing engrams (Espinoza et al., 2018; Johnston et al., 2016; Luna et al., 2019; McAvoy et al., 2016; Sahay et al., 2011b). Consistent with this idea, rAAV-induced ablation of neurogenesis results in a dose-dependent increase in immediate early gene activation 4 weeks after infection (**Fig. 4A,B, S3A**).

In the majority of studies in which rAAV was injected into the DG, neurogenesis was not assessed following viral transduction. However, Danielson et al. (2016) reported *in* vivo calcium imaging of adult-born and mature DGCs using rAAV for delivery of Gcamp6. In this paradigm, abDGCs were labeled via Tamoxifen administration in a Nestin-CreER x tdTomato reporter mouse 3 weeks before rAAV injection into the DG at titers of ~1 e13 gc/mL. Imaging took place 3 weeks later, when tdTomato-labeled cells were ~6 weeks of age. This paradigm would permit tdTomato-labeled abDGCs that were 3 weeks of age at the time of rAAV injection and 6 weeks old at imaging to largely escape rAAV-induced toxicity. However, the loss of abDGCs ~4 weeks old and younger and their contribution to activity in the DG might have been missed.

The retrograde labeling of DGCs by injecting AAV retro into CA3 provides an important advance for future studies of the DG, as DGCs can now be imaged and manipulated using genetic tools while leaving adult neurogenesis intact (Fig 4). Relatively few studies have performed calcium imaging of the DG *in vivo* (Danielson et al., 2016, 2017; Hainmueller and Bartos, 2018; Pilz et al., 2016). Variability in experimental approaches used to deliver genetically encoded calcium indicators to DGCs and inevitable differences in segmentation routines and analysis of calcium traces make direct comparison with these published results difficult. The results described above do not differ substantially with previous findings of ~40-50% of cells being active during recording (Danielson et al., 2016; Pilz et al., 2016) and producing only a few calcium transients per minute (Danielson et al., 2016). However, given the ability of abDGCs to modulate activity within the DG (Ikrar et al., 2013; Lacefield et al., 2012; Luna et al., 2019), viral methods used to deliver calcium indicators and other genetic tools into the DG should be carefully evaluated in future studies.

### Caveats for gene therapy

Based on its stable transgene expression, low risk of insertional mutagenesis, and diminished immunogenicity, rAAV has become the most widely used viral vector for human gene therapy. More than 100 clinical trials using AAV vectors have claimed vector safety (Choudhury et al., 2017; Hocquemiller et al., 2016; Hudry and Vandenberghe, 2019), resulting in two FDA-approved therapies for treating genetic diseases of the CNS (Smalley, 2017, Hoy, 2019). Despite high rates of infection among humans (Thwaite et al., 2015), AAV infection has not been associated with illness or pathology (Büning and Schmidt, 2015). However, recent reports have suggested rAAV may exhibit intrinsic toxicity in multiple tissues (Bockstael et al., 2012; Flotte and Büning, 2018; Hinderer et al., 2018; Hirsch et al., 2011; Hordeaux et al., 2018). Given the steep dose response of rAAV-induced cell death measured in our study, intravenous administration of rAAVs at clinically relevant titers is less likely to cross the blood-brain barrier (BBB) and reach the SGZ of the DG with sufficient MOI to ablate neurogenesis. However, the protective capacity of the BBB does not preclude AAV-induced toxicity in other progenitor and dividing cells throughout the body. Further studies are needed to characterize rAAV-induced toxicity to stem and progenitor cells in other tissues. Also, high MOI may reach the SGZ in CNS studies where rAAV is injected intrathecally (see STRONG trial) or directly into brain tissue (Bartus et al., 2013; Castle et al., 2018; Christine et al., 2009; Hammond et al., 2017; Mandel, 2010; Nagahara et al., 2013; Tuszynski et al., 2015). While attenuation of neurogenesis likely occurs in patients undergoing other treatments such as chemotherapy (Hodge et al., 2008) or radiation treatment (Santarelli et al., 2003; Saxe et al., 2007), the extent of ablation induced by rAAV is striking and shows no signs of recovery throughout the duration of our experiments. This study serves as an additional reminder that rAAV and other viral gene therapies may be associated with significant side effects, particularly during development. Careful consideration of viral titer, delivery method, and viral engineering should be exercised to mitigate side effects where viral therapy may substantially alleviate morbidity or extend life (Mendell et al., 2017).

## METHODS

### Animal use

All animal procedures were approved by the Institutional Animal Care and Use Committees of the Salk Institute and the University of California San Diego, and all experiments were conducted according to the US Public Health Service guidelines for animal research. Wild-type male C57BL/6J mice (Jackson Laboratories), 6 to 7 weeks of age at the time of surgery, were used in this study. Unless otherwise noted, mice were group housed with up to 5 mice per cage in regular cages (RC; 4.7” L × 9.2” W × 5.5” H, InnoVive, San Diego, CA) under standard conditions, on a 12h light–dark cycle, with *ad libitum* access to food and water. BrdU (Sigma) was administered i.p. at 50 mg/kg/day for 3 days.

### Viral injection

Mice were anesthetized with isoflurane (2% via a nose cone, vol/vol), administered with dexamethasone (2.5 mg/kg, i.p.) to decrease inflammation, and placed in a stereotaxic frame. One microliter of virus solution diluted in sterile saline or saline control, unless otherwise specified, was delivered to the hippocampus through stereotaxic surgery using a microinjector (Nanoject III, Drummond Science). Specifically, the difference between bregma and lambda in anteroposterior coordinates was determined. From bregma, DG injection coordinates were calculated as indicated in Table 1.

**Table 1.** DG Injection coordinates. Injection coordinates as measured from bregma adjusted for measured distance between lambda & bregma (Λ-B): anterior-posterior (A/P), medial-lateral (M/L); and dorsoventral depth from dura (D/V).

| Λ-B<br>[mm] | A/P<br>[mm] | M/L<br>[mm] | D/V<br>[mm] |
| --- | --- | --- | --- |
| 3.0 | -1.5 | $\pm 1.5$ | -1.8 |
| 3.2 | -1.6 | $\pm 1.55$ | -1.8 |
| 3.4 | -1.7 | $\pm 1.6$ | -1.9 |
| 3.6 | -1.8 | $\pm 1.65$ | -1.9 |
| 3.8 | -1.9 | $\pm 1.7$ | -1.95 |
| 4.0 | -2.0 | $\pm 1.75$ | -2.0 |

CA3 injection coordinates were calculated as follows: anteroposterior (A/P) −1.8 mm, lateral (M/L) −1.8 mm, ventral (V/L; from dura) −1.6 mm and −2.0 mm, with 400 nL injected at each depth. Following completion of the surgery, carprofen (5 mg/kg, i.p.) and Buprenorphine SR LAB (0.1 mg/kg, s.c.) were administered for inflammation and analgesic relief. Mice were allowed to recover and then returned to their cages. The following viral vectors were used: AAV1-CAG::flex-eGFP-WPRE-bGH (Zeng – Addgene Plasmid #51502, U Penn Vector Core & Addgene), AAVretro-CaMKIIa::NES-jRGECO1a-WPRE-SV40 (Gage, Salk), AAV8-CaMKIIa::NES-jRGECO1a-WPRE-SV40 (Gage, Salk), AAV1-Syn::NES-jRGECO1a-WPRE-SV40 (Kim – Addgene Plasmid# 100854, U Penn), AAV8-CaMKIIa::mCherry-WPRE-bGH (Deisseroth – Addgene Plasmid #114469, Salk), AAV8 capsid (Salk).

### Enriched/novel environments

Mice assigned to enriched environments (EE) were housed in regular caging (RC) and then moved to an EE cage, whereas matched RC controls remained in RC. The EE cage (36” L × 36” W × 12” H) contained a feeder, 2-3 water dispensers, a large and a small running wheel and multiple plastic tubes and domes, and paper huts, with a 12-hour light–dark cycle. Objects in the EE cage were kept constant throughout the experiment; placement of the objects was altered only to the extent that the mice moved them within the cages. Mice were kept in EE or RC for 13 days and injected with BrdU on the final 3 days. On the final day of BrdU, mice were also unilaterally injected with 1 uL 3 e12 gc/mL AAV1-CAG-flexGFP into the DG. Following surgery, animals were returned to EE or RC and sacrificed 2 days post-injection. Mice that received NE exposure for cFOS activation experiments remained in RC until NE exposure and were then transferred to EE cages (as described above) for 15 minutes. Animals were sacrificed and brain tissue collected 1 hour after NE exposure.

### Cranial window placement

For 2-photon calcium imaging experiments, ~1 hour after receiving viral injections as described above, a ~3 mm diameter craniotomy was performed, centered around the DG viral injection site. The underlying dura mater was removed and the cortex and corpus callosum were aspirated with a blunt tip needle attached to a vacuum line. Sterile saline was used to irrigate the lesion and keep it free of blood throughout the surgery. A custom 3-mm diameter, 1.4-mm deep titanium window implant with a 3-mm glass coverslip (Warner Instruments) bottom was placed on the intact alveus of the hippocampus. The implant was held in place with UV-cured dental adhesive (Kerr Dental, Optbond All-In-One) and dental cement (Lang Dental, Ortho-Jet). A small custom titanium head bar was attached to the skull to secure the animal to the microscope stage. Following completion of the surgery, carprofen (5 mg/kg, i.p.) and Buprenorphine SR LAB (0.1 mg/kg, s.c.) were administered (as previously mentioned, animals received 1 dose of each at the end of the final surgery) for inflammation and analgesic relief. Mice were allowed to recover and then returned to their cages. We have previously found that surgical implant and imaging procedures do not affect adult neurogenesis (see Gonçalves et al. 2016, Fig. S2.8 & S9).

### 2-photon calcium imaging of DG activity

Mice were acclimated to head-fixation beginning 1 week after surgery. At time of imaging, each mouse was secured to a goniometer-mounted head-fixation apparatus and a custom-built laser alignment tool was used to level the plane of the cranial window coverslip perpendicular to the imaging path of the microscope objective. Imaging of dorsal DG was performed with a 2-photon laser scanning microscope (MOM, Sutter Instruments) using a 1070 nm femtosecond-pulsed laser (Fidelity 2, Coherent) and a 16× water immersion objective (0.8 NA, Nikon). Images were acquired using the ScanImage software implemented in MATLAB (MathWorks). Imaging sessions were performed intermittently from 10-18dpi to determine optimal viral expression and imaging window. Analyzed activity videos were acquired at ~14dpi in successive 5-minute intervals (512 x 128 pixels; ~3.91Hz).

### Analysis and quantification of calcium activity

Custom software was written in Matlab to extract neuronal activity from 2-photon calcium imaging videos. Calcium traces were extracted by first performing image stabilization for each video using a rigid alignment, maximizing the correlation coefficient between each frame of the movie with an average reference frame constructed from 20-30 frames acquired when the mouse was not running. Alignment was further improved using a line-by-line alignment. Portions of the movie with excessive movement artifact, defined by cross-correlation coefficients below a defined threshold, were discarded. Mice were not trained or rewarded for running on the treadmill and thus were stationary during the majority of the presented calcium imaging data. Automated cell segmentation was achieved by scanning a ring shape of variable thickness and size across a motion corrected reference image. When the cross-correlation metric exceeded a user-adjustable threshold, a circular shaped ROI was generated and the signal extracted. User input was then taken for each video to remove a small number of false positives and labeled cells that evaded automatic classification. Once each cell was labeled and the intensities were recorded, the baseline fluorescence (F) was fit to an exponential curve to eliminate photo-bleaching effects. The change in fluorescence (ΔF) over the baseline fluorescence (F) was then calculated to yield %ΔF/F. Spiking-related calcium events for each cell were defined as fluorescence transients whose amplitude exceeded 7 standard deviations of the negative fluctuations of the %ΔF/F trace. Active cells were defined as having at least 1 spike during a single movie.

### Tissue collection

Mice were deeply anesthetized with ketamine and xylazine (130 mg/kg, 15 mg/kg; i.p.) and perfused transcardially with 0.9% phosphate buffered saline followed by 4% paraformaldehyde (PFA) in 0.1 M phosphate buffer (pH 7.4). Brains were dissected and post-fixed in 4% PFA overnight then equilibrated in 30% sucrose solution.

### Immunohistochemistry

Fixed brains were frozen and sectioned coronally on a sliding microtome at 40-μm thickness, spanning the anterior-posterior extent of the hippocampus, and then stored at −20 °C until staining.

Brain sections were blocked with 0.25% Triton X-100 in TBS with 3% horse serum and incubated with primary antibody in blocking buffer for 3 nights at 4°C. Sections were washed and incubated with fluorophore-conjugated secondary antibodies for 2h at RT. DAPI was applied in TBS wash for 15 min at RT. Sections were washed and mounted with PVA-Dabco or Immu-Mount mounting media. For BrdU staining, brain sections were washed 3× in TBS for 5 minutes, incubated in 2N HCL in a 37°C water bath for 30 minutes, rinsed with 0.1M Borate buffer for 10 minutes at RT, washed 6× in TBS for 5 minutes, and then the above staining procedure was followed.

Primary antibodies used were rat αBrdU (OBT0030, Accurate; NB500-169, Novus; AB6326, Abcam), rabbit αcleaved-CASPASE3 (9661, Cell Signaling), goat αCFOS (sc-52-G, Santa Cruz), rabbit αCFOS (226003, Synaptic Systems), goat αDCX (sc-8066, Santa Cruz), guinea pig αDCX(AB2253, Millipore), chicken αGFAP (AB5541, Millipore), chicken αGFP (GFP-1020, Aves Labs), rabbit αPROX1 (ab101851, Abcam), rabbit αSOX2 (2748, Cell Signaling), rabbit αTBR2 (ab183991, Abcam). Secondary antibodies used were donkey αchicken-AlexaFlour647 (703-605-155), donkey αchicken-AlexaFluor488 (703-545-155), donkey αrat-AlexaFluor647 (712-605-153), donkey αrabbit-Cy5 (711-175-152), donkey αrabbit-Cy3 (711-165-152), donkey αrabbit-AlexaFluor488 (711-545-152) donkey αguinea pig-AlexaFluor488 (706-545-148), donkey αguinea pig-Cy3 (706-165-148), donkey αguinea pig-AlexaFlour647 (706-175-148), donkey αgoat – AlexaFlour647(705-175-147), donkey αgoat – Cy3 (705-165-147), donkey αgoat AlexaFlour488 (705-545-147) – (Jackson Immuno Research Laboratories).

### Histology acquisition and analysis

Images for analysis of neurogenesis and inflammation markers were acquired using a Zeiss laser scanning confocal microscope (LSM 710, LSM 780, or Airyscan 880) using a 20× objective or an Olympus VS-120 virtual slide scanning microscope using a 10× objective. For confocal images (DCX, IBA1, GFAP, SOX2), Z-stacks were obtained through the entirety of the DGC layer, tiles were stitched using Zen software (Zeiss), and images were maximum projected for quantification. Slide scanner images (BrdU, TBR2) were obtained from a single plane. For markers quantified by cell counts (BrdU, TBR2, SOX2, CASPASE3), counting was performed manually. For markers quantified by fluorescent intensity (DCX, IBA1, GFAP), a region of interest was drawn in Zen software, and the average intensity over that region was recorded. For DCX, the region of interest included the full DGC layer and SGZ. Background autofluorescence was corrected by recording the intensity of a neighboring region of CA3 or CA1 devoid of DCX+ cells. For IBA1 and GFAP, the region of interest was the SGZ and hilus, bounded by the inner edge of the granule cell layer and a line drawn between the endpoints of the two blades. No background correction was performed for inflammation markers due to the relatively complete tiling of glia throughout the hippocampus. For each brain, 2-5 images were quantified per side. A blinded observer quantified all images.

Images for analysis of pyknosis and karyorrhexis were obtained on an Airyscan 880 microscope using a 40× objective. Z-stacks were obtained through the entirety of the DGC layer, tiles were stitched using Zen software, and each individual slice of the z-stack was examined. Nuclei were considered abnormal if the DAPI channel showed condensed, uniform labeling throughout the nucleus instead of the typical variation in intensity observed in healthy cells or if nuclei appeared to be fragmenting into uniformly labeled pieces (Bayer and Altman, 1974; Cahill et al., 2017). Two blinded observers quantified these images.

### AAV empty capsid

rAAV8 empty capsids were synthesized and purified using standard CsCl rAAV production protocols by the Salk Viral Core, without the addition of any ITR containing plasmids or sequences.

Electron microscopy quantification was performed at the Salk Institute’s Waitt Advanced Biophotonics Center. 3.5 μL of 3% diluted rAAV empty capsid stock or positive control using viral stock of known concentration (AAV1-CAG-flexGFP) was applied to plasma etched carbon film on 200 mesh copper grids (Ted Pella, 01840-F), 4 grids per stock. Samples were washed 3 times for 5 seconds, stained with 1% Uranyl Acetate for 1 minute, wicked dry with #1 Whatman filter paper, and air dried before TEM exam. For each grid, 4 fields were selected in each of 4 grid squares, for a total of 16 micrographs per grid, 20,000x magnification on a Libra 120kV PLUS EF/TEM (Carl Zeiss), 2kx2k CCD camera. Two blinded observers each quantified all images and the ratio of empty to control virus was calculated.

### Cell culture

Mouse NPCs were obtained from embryonic female C57BL/6 mouse hippocampus and cultured as described previously (Toda et al., 2017). NPCs were cultured in DMEM/F-12 supplemented with N2 and B27 (Invitrogen) in the presence of FGF2 (20 ng/ml), EGF (20 ng/ml), laminin (1 μg/ml) and heparin (5 μg/ml), using poly-ornithine/laminin (Sigma)-coated plastic plates. Medium was changed every 2 days, and NPCs were passaged with Accutase (StemCell Tech) when plates reached confluence.

### *in vitro* rAAV transduction and time lapse imaging

NPCs were seeded onto 96-well plates at a density of 10k cells/well for 24 hours. At 24 hours, medium was changed and supplemented with propidium iodide (1 μg/mL). To serve as baseline, 2 sets of images were acquired 4 hours apart in bright field and red-fluorescence: 5 images per well, 8 wells per treatment, for 5 treatments, on an IncuCyte S3 Live Cell Analysis System (Essen Biosciences, Salk Stem Cell Core and UCSD Human Embryonic Stem Cell Core). AAV1-CAG-flex-eGFP was added with an initial MOI at 1 e7, 1 e6, 1 e5, 1 e4, or H20 control. MOIs were calculated by dividing total viral particles added per well (1 e12, 1 e11, 1 e10, and 1 e9 viral particles, respectively) divided by the initial seeding density of 10k cells/well; the *in vitro* estimate for MOI inflates the number of viral genomes per cell comparted to the *in vivo* estimate for two reasons. First, cells were allowed to proliferate for 28 hours (approximately a 2-3× increase in cells, **Fig. 3B**) before adding virus. Second, *in vitro* viral particles were distributed throughout the growth media and stochastic diffusion was likely to act as a rate-limiting step to viral entry, whereas *in vivo* viral adsorption was more likely to act as the rate-limiting step to viral entry. Images were then acquired every 4 hours for 60 hours. Data were extracted using IncuCyte Analysis software.

### *In vitro* ITR electroporation imaging and FACS analysis

NPCs were collected in nucleofection solution (Amaxa Mouse NSC Nucleofector Kit, Lonza) and electroporated with 5e6 or 1e6 copies/cell of 5’ biotinylated 145bp AAV ITR ssDNA or scrambled control (ITR: 5’-Biotin-AGGAACCCCTAGTGATGGAGTTGGCCACTCCCTCTCTGCGCGCTCGCTCGCTCACTGAGGCCGGGCGACCAAAGGTCGCCCGACGCCCGGGCTTTGCCCGGGCGGCCTCAGTGAGCGAGCGAGCGCGCAGAGAGGGAGTGGCCAA-3’, scramble: 5’-Biotin-CCACATACCGTCTAACGTACGGATTCCGATGCCCAGATATATAGTAGATGTCTTATTTGTGGCGGAATAGCGCCAGAGCGTGTAGGCCAACCTTAGTTCTCCATGGAAGGCATCTACCGAACTCGGTTGCGCGGCCAAATTGGAT-3’, Integrated DNA technologies) or with H20, in triplicate, then plated onto 24-well plates (see schematic in **Fig. 3D**). AAV2 ITR sequences were obtained from NCBI Viral Genome database, NC_001401.2 (Brister et al., 2015). AAV2 ITRs or a sequence largely homologous with the AAV2 ITR sequence were used in the vast majority of rAAV plasmids. Twenty-four-well plates were segregated for imaging and FACS experiments. Imaging plates were supplemented with propidium iodide and images were acquired on an IncuCyte S3 Live Cell Analysis System as follows: 16 images per well, 4 wells per replicate, 3 replicates per treatment, for 5 treatments, every 3 hours for 60 hours. Data were extracted using Incucyte Analysis software using the same mask definition obtained above. NPCs from FACS plates were collected in PBS at 12, 24, and 36 hours using Accutase. After incubation for 30 minutes at RT with Vybrant DyeCycle Green Stain (ThermoFischer, 1:2000), Zombie UV Fixable Viability Kit (BioLegend, 1:1000) and CountBright Absolute Counting Beads (~5000 beads/sample, Thermofisher), cells were filtered into polypropylene FACS collection tubes and FACS analysis was performed on an LSRFortessa X-20 (BD Biosciences, UCSD Human Embryonic Stem Cell Core). Samples were collected by gating on 1000 CountBright Counting Bead counts per well, 1 well per replicate, 3 replicates per treatment, for the 5 treatments, interleaved, at 3 time points. Populations of live and dead cells (UV Zombie negative and positive cells, respectively), and G1-phase and replicating (S- & G2-phase) cells (Vybrant DyeCycle Green low and high, respectively) were determined using FlowJo software.

### Statistical analysis

All data are presented as mean ± s.d. when describing data between individual samples, and as mean ± s.e.m when comparing between groups. To compare histology data across experiments, counts and intensity measures for the injected side of the DG are presented as a percentage of that group’s mean counts or intensity on the control side. Statistical comparisons were performed in Prism 8.0 (GraphPad Software) using paired t-test (paired data, one independent variable: treatment), repeated measures 2-way ANOVA using either the Tukey or Sidak multiple comparison test (two independent variables: treatment and time) or 2-way ANOVA using Dunnett’s or Tukey’s multiple comparison test (*in vitro* rAAV transduction relative to H2O control and electroporation, respectively) when interaction was significant. Linear regression was performed for BrdU vs cFOS activation. K-S tests were performed for cumulative distributions. All statistical tests were two-tailed. Threshold for significance (α) was set at 0.05; * is defined as p<0.05, ** is defined as p<0.01, *** is defined as p<0.001, n.s. is not significant.

## Acknowledgments

We thank Dr. Matt Hirsch, Dr. Jude Samulski, Dr. Tomo Toda, Dr. Simon Schafer, Dr. Carol Marchetto, Lynne Moore, Ruth Keithley for experimental advice, and Mary Lynn Gage and Dr. Marianna Alperin for comments on the manuscript. This work was made possible by our funding sources: NIH R01 MH090258, NIH R01 MH095741 NIH R01 AG056306, NIH K08 NS093130, The McKnight Endowment Fund for Neuroscience, JPB Foundation, Annette Merle-Smith, James S. McDonnell Foundation, Mathers Foundation, the Leona M. and Harry B. Helmsley Charitable Trust grant # 2012-PG-MED002, Ray and Dagmar Dolby Family Fund, Salk Innovation Grant, and the Dan and Martina Lewis Biophotonics Fellows Program. Special thanks to Salk Cores: Waitt Advanced Biophotonics Core with funding from NIH-NCI CCSG: P30 014195 and the Waitt Foundation, STEM Core, Viral Vector Core supported by Salk Cancer Center, NCI P30 CA014195; and UCSD Human Embryonic Stem Cell Core.

