## supplement for "AAV Ablates Neurogenesis in the Adult Murine Hippocampus"

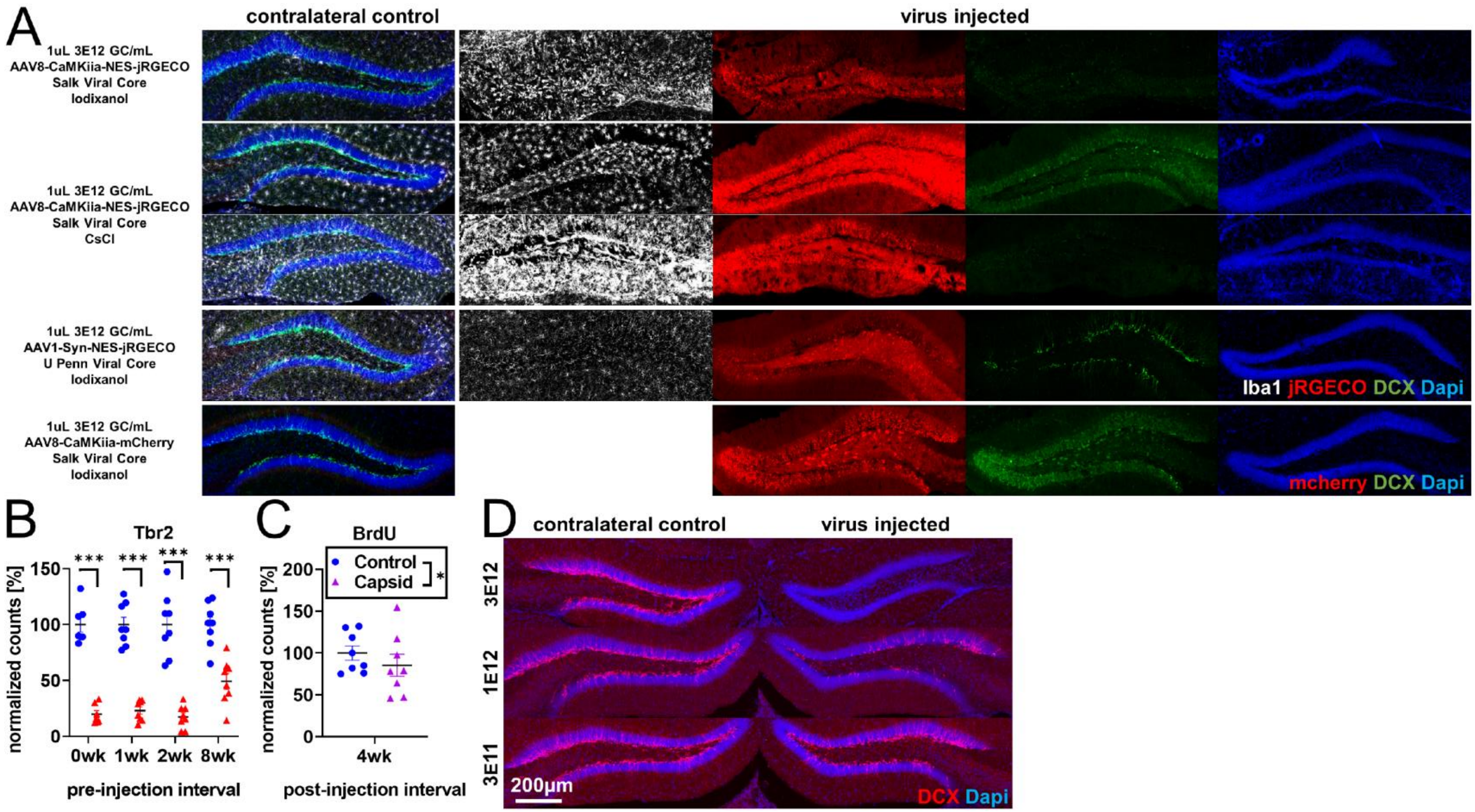


**Figure S1 rAAV-dependent toxicity (A)** rAAV-induced reduction of DCX expression is variable approximately 2-weeks post viral injection, but independent of viral core, purification method, serotype, promoter, and protein expressed. Examples in rows 2 and 3 are taken from different animals injected with the same titer and volume of AAV8-CaMKiia-NES-jRGECO (Salk Viral Core – CsCl purification), demonstrating variability in magnitude, but consistency in direction of Iba1+ microglia activation and loss of immature neuronal marker DCX staining across animals. **(B)** In the same tissue as Figure 1 A-B, Tbr2+ intermediate progenitors are lost even when older BrdU+ cells are spared. **(C)** Consistent with Figure 1B last column, BrdU+ cells are intact 4 weeks after empty capsid injection. **(D)** Consistent with Figure 1 C & D, injected titers of 3 E12 result in nearly complete ablation of DCX expression, injected titers of 1 E12 result in partial abolition of DCX expression, and injected titers of 3 E11 result in a small loss of DCX expression at 4 weeks post virus. Data are presented as mean ± s.e.m, significance reported as: *** $p < 0.001$.

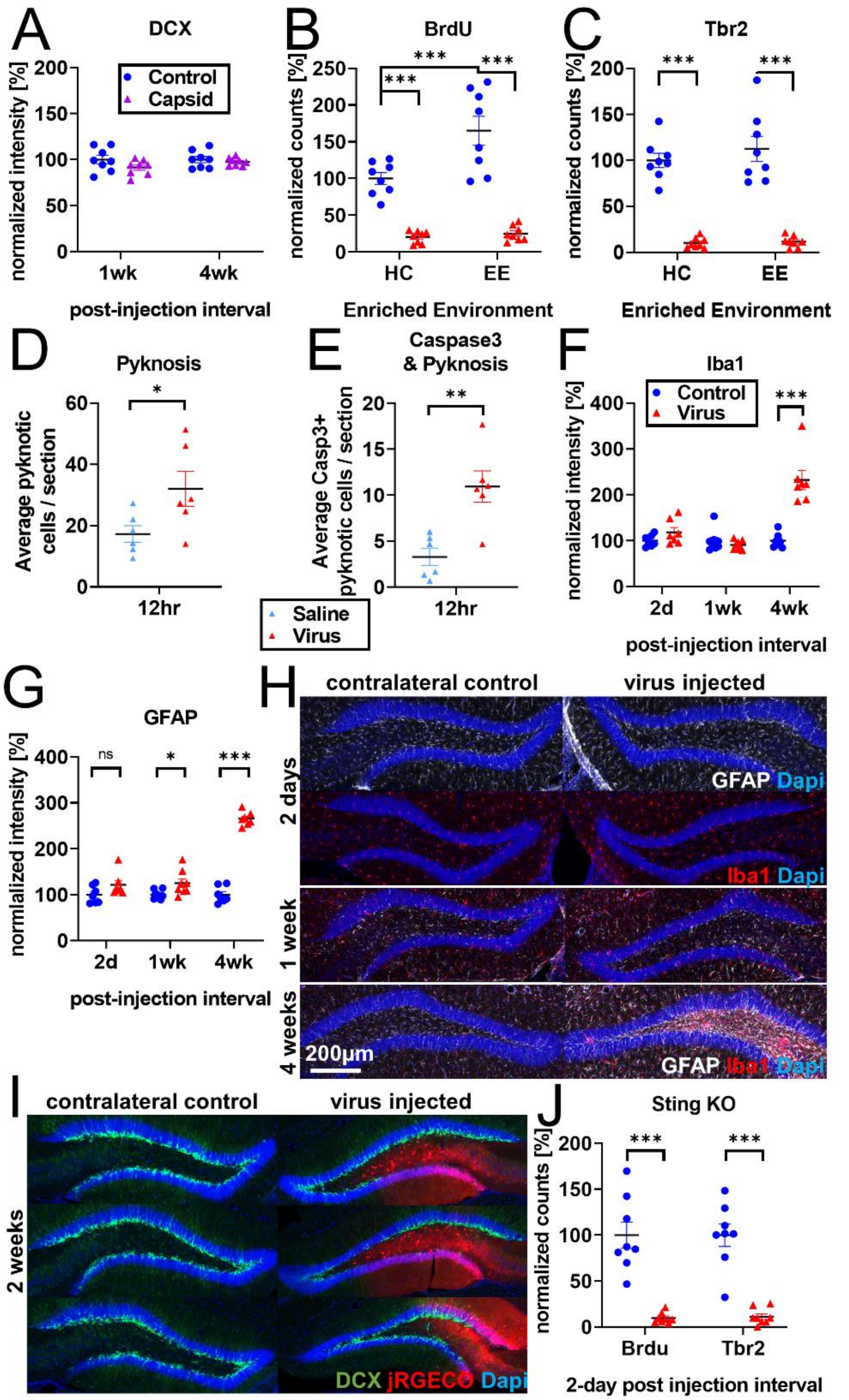


**Figure S2. Inflammation and cell loss (A)** Consistent with Figure 1B, S1C, 2F, immature neuronal marker DCX is preserved at 1 and 4 weeks post-injection of empty capsid. **(B)** Environmental enrichment is sufficient to increase adult neurogenesis and increase BrdU labeling but insufficient to protect against rAAV-induced cell loss. **(C)** Enriched environment is insufficient to protect Tbr2+ intermediate progenitors against rAAV induced cell loss**. (D)** Pyknotic cells in the granule cell layer and SGZ show a significant increase in rAAV-injected DGs 12 hours post-injection relative to saline control. **(E)** Casp3+ pyknotic cells in the granule cell layer and SGZ are significantly increased 12 hours post-rAAV injection relative to saline controls. **(F)** Extending experiments from Figure 2A-E, Iba1 intensity is not significantly changed in subgranular zone (SGZ) and hilus 2 days following rAAV injection and greatly increased 4 weeks following injection. **(G)** GFAP intensity is slightly increased in SGZ and hilus 1 week following rAAV injection and greatly increased 4 weeks following injection. **(H)** Representative images of Iba1 and GFAP histology; increases in Iba1 and GFAP activation are not commensurate with rAAV-induced toxicity. No obvious change in microglial morphology was observed at 2 days or 1 week relative to contralateral control; at 4 weeks microglia exhibit activated amoeboid morphology. **(I)** Following 30-nL injections of 5E12 GC/mL AAV1-Syn-NES-jRGECO into DG, loss of DCX labeling is incomplete and follows boundaries of viral spread. **(J)** Injection of 1 µL 5E12 GC/mL AAV1-Syn-NES-jRGECO into Sting-KO mice results in BrdU+ and Tbr2+ cell loss. All data are presented as mean ± s.e.m, significance reported as: * $p < 0.05$, ** $p < 0.01$, *** $p < 0.001$.

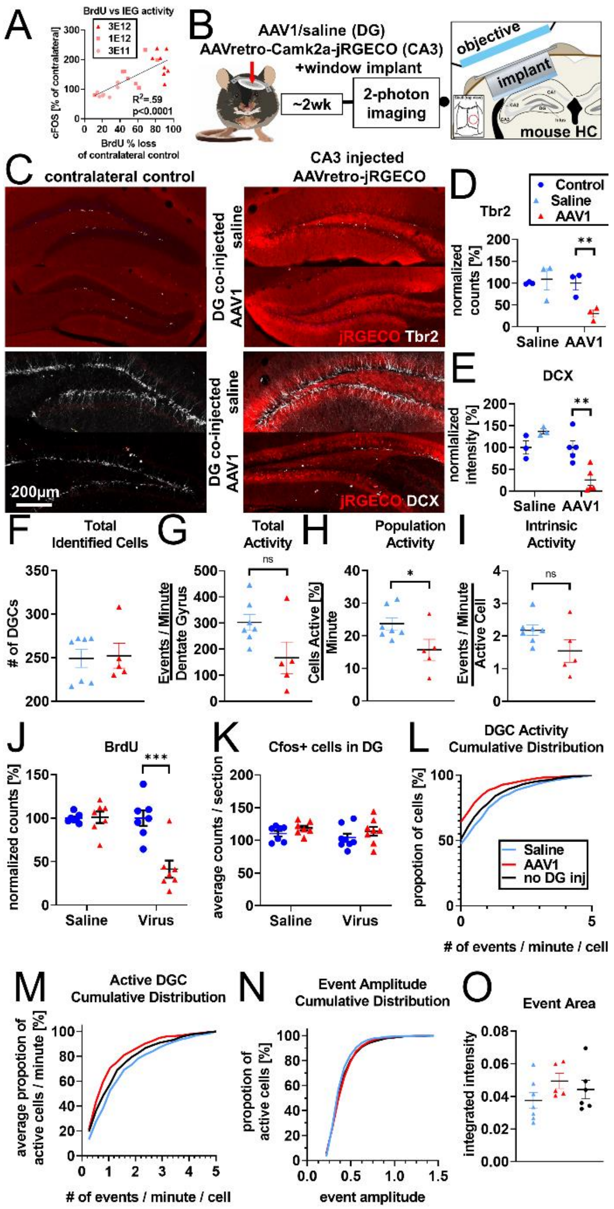


**Figure S3. Calcium imaging following 2-week knockdown of adult neurogenesis. (A)** Additional analysis of experiments described in Figure 1E demonstrate that cFOS activation in mature DGCs is significantly correlated with knockdown efficiency of adult neurogenesis**. (B)** Experimental design for 2-photon imaging of DG utilizing AAVretro. 800 nL of 3 E12 gc/mL AAVretro-CaMKiia-jRGECO is injected into CA3 and either 1µL 3 E12 AAV1-CAG-flex-eGFP or saline is injected into the DG. Mice are implanted with a cranial window and undergo 2-photon calcium imaging 2 weeks later. **(C)** Representative images show Tbr2+ and DCX+ cells are intact in animals injected with AAV retro in CA3 when saline is injected into DG but not when AAV1 is injected into DG. **(D)** Quantification of Tbr2+ cells demonstrates adult neurogenesis is intact in saline co-injected animals but not in AAV1 co-injected animals ($F_{side\ x\ treatment}$ (1,4)=8.3, p<0.05; saline: 8.8% ± 25.8%, n.s.; AAV1: -69.5% ± 8.3±, p<0.01). **(E)** Quantification of DCX staining demonstrates adult neurogenesis is intact in saline co-injected but not in AAV1 co-injected animals ($F_{side\ x\ treatment}$ (1,6)=11.8, p<0.05; saline: 36.80% ± 20.6, n.s.;AAV1:-74.40% ± 21.5%, p<0.01). **(F)** Total number of identified DGCs in each animal is not different between treatment groups (3.1 ± 17.1 cells, t(10)=0.2, n.s.). **(G)** AAV1 co-injected animals show a trend toward having fewer total spontaneous calcium transients 2 weeks after injection than saline controls (-135.3 ± 62.4 events/minute/DG, t(10)=2.2, n.s.). **(H)** This effect is more apparent in the number of active cells of AAV1 co-injected animals (-8.0 ± 3.5 cell active [%]/minute, t(10)=2.3, p<0.05) but does not reach significance in **(I)** the level of spontaneous activity per individual active cells (-0.6 ± 0.3 events/minute/active cell, t(10)=1.8, n.s.). **(J)** A mock-imaging experiment to compare calcium imaging results with IEG activity. Animals underwent the same procedure as above but were placed in novel environment instead of undergoing calcium imaging. Adult neurogenesis is intact in saline-injected animals but not in AAV1 co-injected animals (($F_{side\ x\ treatment}$ (1,12)=12.9, p<0.01; saline: 1.0% ± 7.8%, n.s.;AAV1:-58.6% ± 14.7%, p <0.001). **(K)** However, there is no difference in number of cfos+ cells in the DG ($F_{side}$ (1,13)=2.3, n.s.; $F_{treatment}$ (1,13)=1.1, n.s.; $F_{side\ x\ treatment}$ (1,13)=0.01 n.s.;) at 2 weeks post viral injection. **(L)** Comparing across the AAV retro imaging experiment with no direct viral injection into the DG (**Figure 4**) and 2-week knockdown experiment above, the increase in inactive cells seen with co-injection of AAV1 into the DG (red) can be seen in a left-shifted cumulative distribution, where saline co-injected animals and animals that received no DG injection are highly similar. **(M)** The cumulative distribution of active cells demonstrates little difference in the firing rate of active DGCs between conditions. **(N)** The distribution of DGC event amplitudes is similar between treatment groups. **(O)** Size of DGC events is not different between treatments ($F_{treatment}$ (2,15)=1.3, n.s.). All data are presented as mean ± s.e.m, significance reported as: * p < 0.05, ** p < 0.01, *** p < 0.001.

**Supplementary Table 1. Tukey's test for group comparisons of AAV ITR and SCR control.** Post hoc comparisons of the electroporation groups presented in Fig 3E and 3F following 2-way ANOVA. * $p < 0.05$, ** $p < 0.01$, *** $p < 0.001$, n.s. = not significant.

| Cell growth (% Confluence) | | | | | | | | | | |
|---|---|---|---|---|---|---|---|---|---|---|
| Time (min) | H20 vs. SCR 1E6 | H20 vs. SCR 5E6 | H20 vs. ITR 1E6 | H20 vs. ITR 5E6 | SCR 5E6 vs. SCR 1E6 | ITR 1E6 vs. SCR 1E6 | ITR 5E6 vs. SCR 1E6 | ITR 1E6 vs. SCR 5E6 | ITR 5E6 vs. SCR 5E6 | ITR 5E6 vs. ITR 1E6 |
| 3 | n.s. | n.s. | n.s. | n.s. | n.s. | n.s. | n.s. | n.s. | n.s. | n.s. |
| 6 | n.s. | n.s. | n.s. | * | n.s. | n.s. | * | n.s. | n.s. | ** |
| 9 | n.s. | n.s. | n.s. | *** | n.s. | n.s. | ** | n.s. | * | *** |
| 12 | n.s. | n.s. | n.s. | *** | n.s. | n.s. | *** | n.s. | ** | *** |
| 15 | n.s. | n.s. | n.s. | *** | n.s. | n.s. | *** | n.s. | *** | *** |
| 18 | n.s. | n.s. | n.s. | *** | n.s. | n.s. | *** | n.s. | *** | *** |
| 21 | n.s. | n.s. | n.s. | *** | n.s. | n.s. | *** | * | *** | *** |
| 24 | n.s. | n.s. | n.s. | *** | n.s. | n.s. | *** | ** | *** | *** |
| 27 | n.s. | * | n.s. | *** | n.s. | n.s. | *** | ** | *** | *** |
| 30 | n.s. | ** | n.s. | *** | n.s. | n.s. | *** | *** | *** | *** |
| 33 | n.s. | ** | n.s. | *** | * | n.s. | *** | *** | *** | *** |
| 36 | n.s. | ** | n.s. | *** | * | n.s. | *** | *** | *** | *** |
| 39 | n.s. | * | n.s. | *** | ** | n.s. | *** | *** | *** | *** |
| 42 | n.s. | * | n.s. | *** | * | n.s. | *** | ** | *** | *** |
| 45 | n.s. | * | n.s. | *** | n.s. | n.s. | *** | n.s. | *** | *** |
| 48 | n.s. | *** | n.s. | *** | n.s. | n.s. | *** | n.s. | *** | *** |
| 51 | n.s. | ** | n.s. | *** | n.s. | n.s. | *** | n.s. | *** | *** |
| 54 | n.s. | n.s. | n.s. | *** | n.s. | n.s. | *** | n.s. | *** | *** |
| 57 | n.s. | n.s. | n.s. | *** | n.s. | n.s. | *** | n.s. | *** | *** |
| 60 | n.s. | n.s. | n.s. | *** | n.s. | n.s. | *** | n.s. | *** | *** |
| Cell death (Proportion Propidium Iodide+) | | | | | | | | | | |
| Time (min) | H20 vs. SCR 1E6 | H20 vs. SCR 5E6 | H20 vs. ITR 1E6 | H20 vs. ITR 5E6 | SCR 5E6 vs. SCR 1E6 | ITR 1E6 vs. SCR 1E6 | ITR 5E6 vs. SCR 1E6 | ITR 1E6 vs. SCR 5E6 | ITR 5E6 vs. SCR 5E6 | ITR 5E6 vs. ITR 1E6 |
| 3 | n.s. | n.s. | n.s. | *** | n.s. | n.s. | *** | n.s. | *** | *** |
| 6 | ** | *** | *** | *** | n.s. | * | *** | n.s. | *** | *** |
| 9 | *** | *** | *** | *** | n.s. | ** | *** | * | *** | *** |
| 12 | *** | *** | *** | *** | n.s. | * | *** | n.s. | *** | *** |
| 15 | *** | *** | *** | *** | n.s. | * | *** | ** | *** | *** |
| 18 | *** | *** | *** | *** | n.s. | n.s. | *** | n.s. | *** | *** |
| 21 | ** | *** | *** | *** | n.s. | n.s. | *** | n.s. | *** | *** |
| 24 | * | *** | *** | *** | n.s. | n.s. | *** | n.s. | *** | *** |
| 27 | n.s. | *** | *** | *** | n.s. | n.s. | *** | n.s. | *** | *** |
| 30 | n.s. | ** | *** | *** | n.s. | n.s. | *** | n.s. | *** | *** |
| 33 | n.s. | ** | *** | *** | n.s. | n.s. | *** | n.s. | *** | *** |
| 36 | n.s. | * | ** | *** | n.s. | n.s. | *** | n.s. | *** | *** |
| 39 | n.s. | n.s. | ** | *** | n.s. | * | *** | n.s. | *** | *** |
| 42 | n.s. | n.s. | *** | *** | n.s. | * | *** | n.s. | *** | *** |
| 45 | n.s. | n.s. | *** | *** | n.s. | * | *** | * | *** | *** |
| 48 | * | n.s. | *** | *** | n.s. | n.s. | *** | ** | *** | *** |
| 51 | ** | n.s. | *** | *** | n.s. | n.s. | *** | *** | *** | *** |
| 54 | * | n.s. | *** | *** | n.s. | n.s. | *** | *** | *** | *** |
| 57 | * | n.s. | *** | *** | n.s. | n.s. | *** | *** | *** | *** |
| 60 | n.s. | n.s. | *** | *** | n.s. | n.s. | *** | ** | *** | *** |
